# Interplay between meropenem and human serum albumin on expression of carbapenem resistance genes and natural competence in *Acinetobacter baumannii*

**DOI:** 10.1101/2021.05.14.444273

**Authors:** Casin Le, Camila Pimentel, Marisel R. Tuttobene, Tomas Subils, Brent Nishimura, German M. Traglia, Federico Perez, Krisztina M. Papp-Wallace, Robert A. Bonomo, Marcelo E. Tolmasky, Maria Soledad Ramirez

## Abstract

*Acinetobacter baumannii* A118, a mostly susceptible strain, and AB5075, carbapenem-resistant, were cultured in L broth or L broth with different supplements: 3.5% human serum albumin (HSA), human serum (HS), meropenem, or meropenem plus 3.5% HSA. Natural transformation levels were enhanced in *A. baumannii* A118 and AB5075 cultured in medium supplemented with 3.5 % HSA. Addition of meropenem plus 3.5% HSA caused synergistic enhancement of natural transformation in *A. baumannii* A118. Medium containing 3.5% HSA or meropenem enhanced the expression levels of the competence and type IV pilus associated genes. The combination meropenem plus 3.5% HSA produced a synergistic augmentation in the expression levels of many of these genes. The addition of HS, which has a high content of HSA, was also an inducer of these genes. Cultures in medium supplemented with HS or 3.5% HSA also affected resistance genes, which were expressed at higher or lower levels depending on the modification required to enhance resistance. The inducing or repressing activity of these modulators also occurred in three more carbapenem-resistant strains tested. An exception was the *A. baumannii* AMA16 *bla*_NDM-1_ gene, which was repressed in the presence of 3.5% HSA. In conclusion, HSA produces an enhancement of natural transformation and a modification in expression levels of competence genes and antibiotic resistance. Furthermore, when HSA is combined with carbapenems, which may produce stronger cellular stress, the *A. baumannii* responds increasing the levels of expression of genes involved in natural competence. This process may favor the acquisition of foreign DNA and accelerate evolution.

**Importance:** *Acinetobacter baumannii* causes a variety of nosocomial- and community-infections that are usually resistant to multiple antimicrobial agents. As new strains acquire more resistance genes, these infections become harder to treat, and mortality can reach up to 39%. The high genomic plasticity exhibited by *A. baumannii* must be the consequence of numerous mechanisms that include acquiring foreign DNA and recombination. Here, we describe the ability of *A. baumannii* to induce competence genes when exposed to environments that resemble those found in the human body during untreated infection or after administration of carbapenems. In this latter scenario expression of genes related to resistance also modify their expression levels such that resistance is increased. The contributions of this article are two-pronged. First, when *A. baumannii* is exposed to substances present during infection, it responds, augmenting the ability to capture DNA and accelerate evolution. Second, in those conditions, the bacterium also modifies the expression of resistance genes to increase its resistance levels. In summary, recognition of substances that are naturally (HSA) or artificially (treatment with carbapenems) induces *A. baumannii* to defend, enhancing resistance and increasing the chances of acquiring new resistance mechanisms.

## 1. Introduction

Due to increased prevalence and the lack of effective antibiotics and antibiotics in development to target carbapenem-resistant *A. baumannii*, the Centers for Disease Control and Prevention has classified it as an urgent threat to human health (1, 2). Alarmingly, most carbapenem-resistant *A. baumannii* are extensively-drug resistant with little to no treatment options available (3). The prevalence of carbapenem-resistant *A. baumannii* has increased around the World (40-80% of isolates are carbapenem-resistant) with infection control measures being one of the few effective strategies to prevent spread (3-5). In USA, carbapenem-resistant *A. baumannii* rates in central line-associated bloodstream and catheter-associated urinary tract infections reach 47 and 64%, respectively (3). In the year 2017, Europe reported that 33.4 % of the bloodstream infections caused by *A. baumannii* were carbapenem-resistant and in some countries, particularly those in Southern and Eastern Europe, carbapenem resistance reaches 80% (3). Among the properties that have permitted *A. baumannii* to evolve and persist in clinical settings is its tremendous genetic plasticity driven by its capacity to acquire foreign DNA, especially antimicrobial resistance genes. The genomes of *A. baumannii* are highly variable and include large DNA segments from different origins, thus highlighting the role of natural transformation in its rapid evolution and acquisition of survival traits (6-18).

Natural transformation, a process described and characterized in more than 80 species of Gram-positive and Gram-negative bacteria, is the ability of competent bacteria to take up exogenous DNA and integrate it into their chromosome (19-23). Bacteria require specific stimuli to develop natural competence (24-26). For example, the major protein component of blood, human serum albumin (HSA) enhances transformation frequency and increases competence genes expression in *A. baumannii* strains (11, 13). Human pleural and ascites fluid, whole blood, and human serum, which contain HSA at different concentrations, also enhance DNA acquisition (13, 18). Most often patients acquire bloodstream infections due to *A. baumannii* secondary to pneumonia (4). Other risk factors for acquiring *A. baumannii* bacteremia include a urinary tract infection, contaminated healthcare equipment as well as a surgical site infection. Mortality risk ranges from 20-39% although comorbidities as well as bacteremia due to multi-drug resistant strains of *A. baumannii* further increases mortality (4). Thus, the ability of *A. baumannii* to modify its genome within the human host and in patients with bacteremia through the acquisition of foreign DNA is alarming. These processes illustrate the intricate link between the host-pathogen interactions and an enhancement in the capacity of *A. baumannii* to evolve through gene acquisition. Furthermore, antibiotics that are or were used to treat *A. baumannii* infections, such as mitomycin, kanamycin, streptomycin, meropenem, and quinolones further increase competence and facilitate the acquisition of resistance genes (12, 27, 28). These antibiotics were or are currently used as treatment options for infections caused by *A. baumannii*; thus, these antimicrobials can further facilitate the acquisition of resistance genes (29).

Several studies suggest a correlation between natural transformation and the type IV pilus-dependent movement of *A. baumannii* on surfaces (6, 10, 30). *A. baumannii*’*s* type IV pilus is an essential component of the DNA-uptake machinery (24). The type IV pilus is composed by 21 proteins, where PilA is the major pilin subunits. Among them, there are also ATPases such PilB and PilT/PilU that mediate the extension and retraction process (30).The DNA enters the periplasmic space through the PilQ secretin, where it is bound by the DNA-binding protein ComEA. DNA translocates through the ComEC channel and, as in most cases of natural transformation, one of the strands is degraded. Once in the cytosol, Ssb, DrpA, RecA and ComM are proteins responsible for DNA binding and recombination. Expression of the type IV pilus is regulated by two two-components systems, the PilRS and the Pil-Chp chemosensory system (30-34). In 2021, Vesel and Blokesch demonstrated that the transformability of *A. baumannii* A118 strain varied significantly during different growth phases. This variability was likely due to growth phase-dependent production of its type IV pilus; moreover, pilus-related genes were essential for the bacterium’s transformability and its surface motility (30).

We assessed the role of HSA alone and in combination with meropenem on enhancing the natural competence of *A. baumannii* by using a concentration range found in human blood (3.4 - 5.4 %). The presence of HSA was associated with increased expression of competence genes and ability to take up foreign DNA in *A. baumannii* strains A118 (a carbapenem-susceptible strain) and AB5075 (a carbapenem-resistant strain). Surprisingly, a synergistic increase in natural transformation and expression of competence genes was also observed combining HSA and meropenem for the carbapenem-susceptible A118 strain, while for the carbapenem-resistant strain only the synergy was observed in the expression of competence genes. Furthermore, the expression of genes associated with carbapenem resistance were modified in presence of HSA at a physiological concentration. In addition, when cells were cultured in human serum (HS), an increased in the expression of natural transformation and carbapenem-resistance genes were further confirmed. Our results showed an interplay between HSA with carbapenems, augmenting the capacity of *A. baumannii’s* DNA acquisition and expression of carbapenem-resistant genes.

## 2. Results and Discussion

To describe in more detail the role of HSA on the expression of competence-associated genes leading to natural transformation, we must briefly review previous transcriptomic data obtained by our lab (35-38). Specifically, data previously obtained for *A. baumannii* A118 and AB5075 cultured in the presence of HSA or human fluids with different amounts of HSA, such as human pleural fluids (HPF) and cerebrospinal fluid (CSF), was analyzed. Based on the Vesel *et al*. publication (30), forty-one competence and type IV pilus associated genes were selected for further examination. (30). These genes participate in type IV pilus synthesis and regulation, DNA uptake and translocation, and ssDNA binding and recombination (Fig. S1 and Table S1). The expression of 30 competence and type IV pilus associated genes was enhanced when *A. baumannii* A118 was cultured in medium supplemented with HSA or HPF. When the supplement was CSF, the number of genes up-regulated was 25 (Table 1, Fig. S1 and Table S1). RNA-seq data indicated that, in the case of *A. baumannii* AB5075, 11, 27, and 18 competence and type IV pilus associated genes were up-regulated in cultures supplemented with HSA, HPF and CSF, respectively (Table 1, Fig. S1 and Table S1).

**Table 1.**
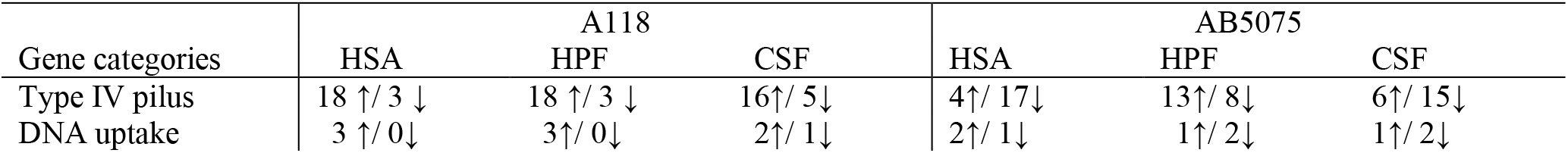
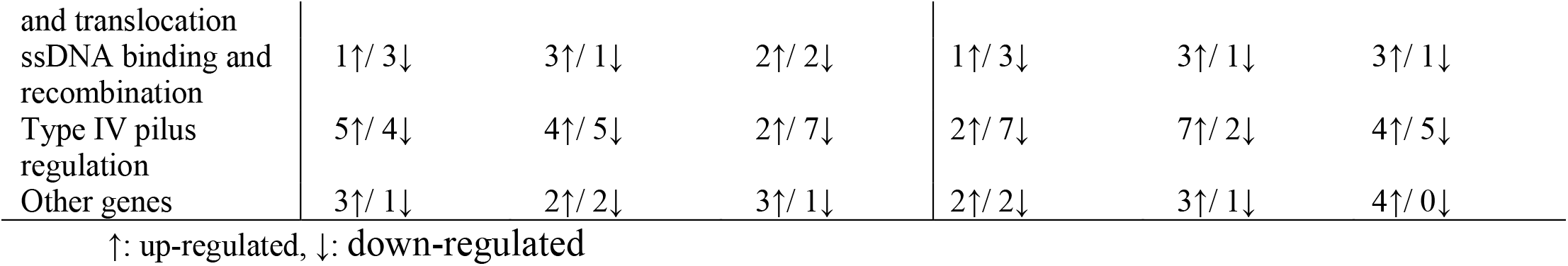
Transcriptomic analysis of type IV pilus associated genes in *A. baumannii* A118 and AB5075 strains, exposed to 0.2% HSA, 4% HPF and 4% CSF, showing the up- and down-regulated genes (35, 36, 38).

### 2.1. Physiological concentrations of HSA enhances both transformation frequencies and expression of competence-related genes

HSA is the most abundant protein in human blood, which represents the 50/60% of the total protein content (13, 39). During the course of an infection, *A. baumannii* is exposed to ∼ 3.4-5.4% HSA. We had observed that human fluids with different content of HSA and 0.2% human or bovine albumins enhanced DNA acquisition and triggered the expression of *comEA* and *pilQ*, herein the effect of the concentration of HSA normally encountered in human blood by *A. baumannii* was evaluated. *A. baumannii* A118 and AB5075 showed a significant increase (*P-value*: 0.0489 and 0.0111, respectively) in transformation frequency, 4.2 and 11-fold-change, respectively, when cultured in medium supplemented with 3.5% HSA (Fig. 1A). The expression levels of the competence and type IV pilus associated genes *pilA, pilT, pilQ, comEA, comEC, comF and drpA* were also increased in these growth conditions (Figure 1 B). The levels of enhancement of expression of these genes in *A. baumannii* A118 and AB5075 were as follows: *pilA*, 16-fold and 4.6-fold; *pilQ*, 5.1-fold and 2.4-fold; *pil*T, 3.6-fold and 2.8-fold; *comEA* 2.8-fold and 1.3-fold; *comEC*, 25-fold and 3.3-fold; *comF*, 23.8-fold and 3.9-fold; and *drpA*, 25-fold and 2-fold. These results are consistent with the increase of natural transformation observed in the presence of physiological concentrations of HSA.

**Figure 1.**
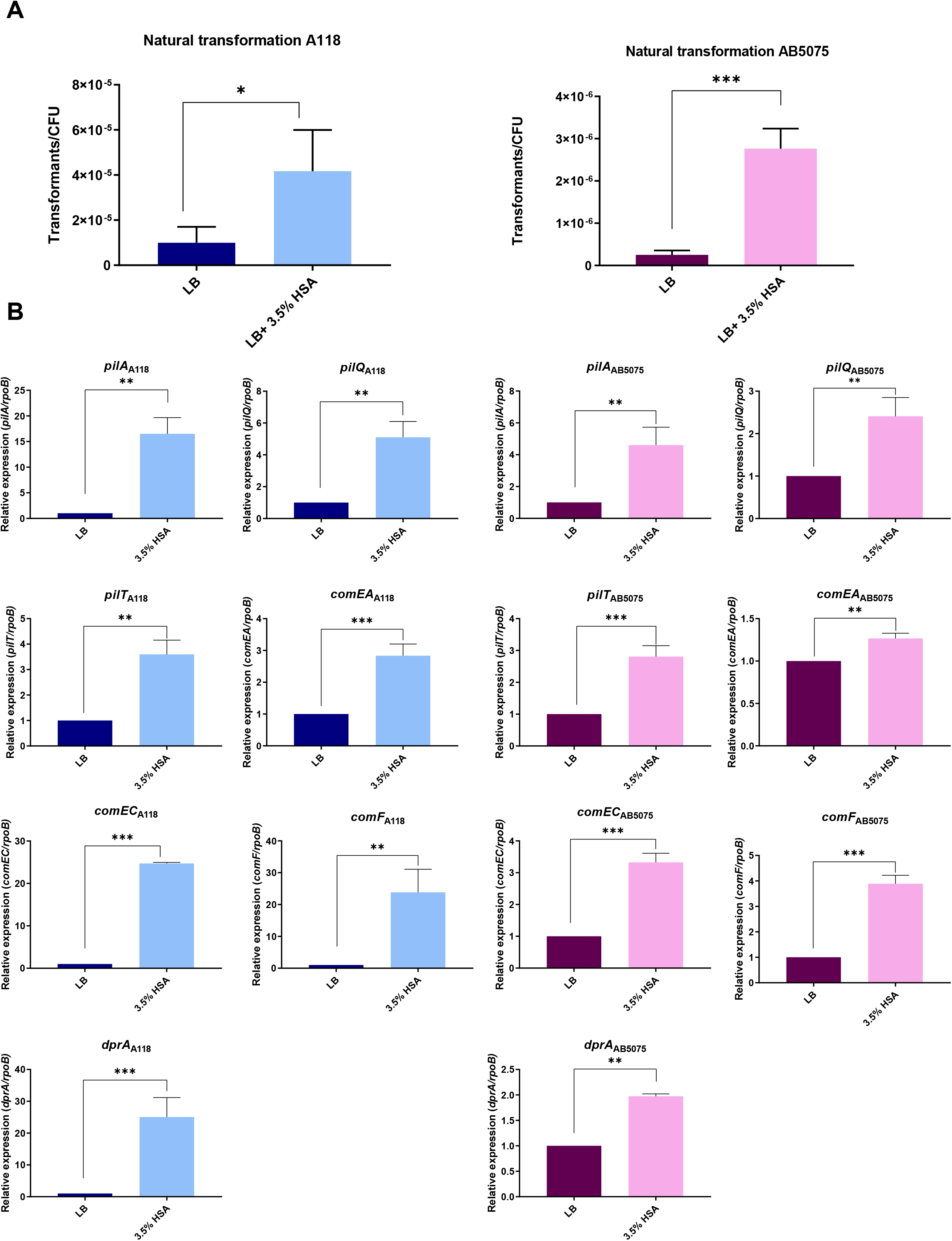
A) Natural transformation frequencies with LB broth or LB broth plus 3.5 % HSA for A118 and AB5075 *A. baumannii* strains. At least three independent replicates were performed and *P-value* < 0.05 was considered significant (*t test*). B) qRT-PCR of A118 and AB5075 strains genes associated with competence and type IV pilus, *pilA, pilQ, pilT, comEA, comEC, comF* and *dprA* in the presence of 3.5% HSA. Fold changes were calculated using double ΔCt analysis. At least three independent samples were used, and three technical replicates were performed from each sample. Statistical significance (*P* < 0.05) was determined by ANOVA followed by *t* test; one asterisks: *P* < 0.05; two asterisks: *P* < 0.01 and three asterisks: *P* < 0.001.

### 2.2. Carbapenem-resistance associated genes expression is modified by the presence of HSA at the concentration range found in blood

Our previous results showed that addition of 0.2% HSA to *A. baumannii* A118 culture medium was correlated with enhanced expression of *bla*_OXA-51-like_, reduced expression of *carO*, and 0.65-fold increase in the MIC of meropenem (38).

Meropenem and imipenem are the carbapenems of choice to treat bloodstream infections caused by carbapenem-susceptible *A. baumannii* (40, 41). Considering the increased resistance to one of these antibiotics observed in cells growing in the presence of 0.2% HSA, it became important to determine if the blood concentrations of HSA had a bigger impact in the levels of resistance. Addition of 3.5% HSA to the culture medium was associated to increased expression of *bla*_OXA-51-like_ in *A. baumannii* A118 and AB5075 (1.44-fold and 3.63-fold, respectively). This condition was also associated to a reduction in expression levels of *carO* (Fig. 2A). Also, addition of 3.5% HSA to the culture medium was also correlated with a 30-fold increase in expression of the *A. baumannii* AB5075 *bla*_OXA-23-like_.

**Figure 2.**
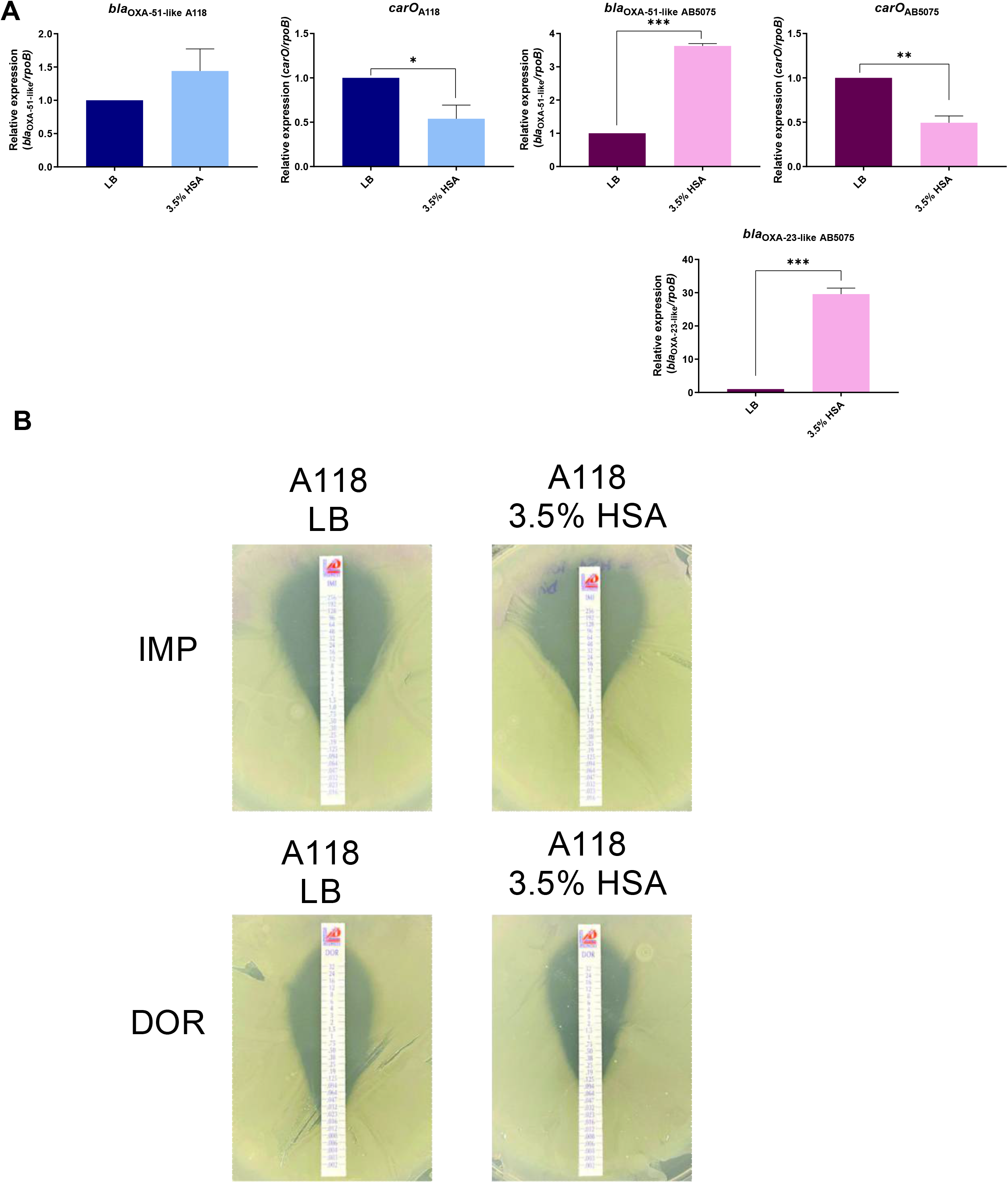
qRT-PCR of carbapenem resistance related genes (*bla*_OXA-51-like_, *bla*_OXA-23-like_ and *carO*) for the *A. baumannii* A118 and AB5075 strains in the presence of 3.5% HSA. Bacterial cells grown in LB and LB plus 3.5% HSA were used. Fold changes were calculated using double ΔCt analysis. At least three independent samples were used, and three technical replicates were performed from each sample. Statistical significance (*P* < 0.05) was determined by ANOVA followed by *t* test, one asterisks: *P* < 0.05; two asterisks: *P* < 0.01 and three asterisks: *P* < 0.001. B) Strain A118 grew in LB broth or LB broth plus 3.5% HSA were used to performed imipenem (IMP) and doripenem (DOR) susceptibility. Minimum inhibitory concentration (MIC) was performed by E-test (Liofilchem, Italy) following CLSI recommendations.

*A. baumannii* A118 cultures in the presence of 3.5 % HSA produced a 2.5- and 1.5-fold change increase in the MIC values of imipenem and doripenem, respectively (Fig. 2B and Table S2). Changes in MICs were not observed for the A118 strain with meropenem and ertapenem between 3.5% HSA and the control (Table S2). The observed increase in resistance towards imipenem and doripenem supports the observed changes in expression for the carbapenem-resistance genes as well as the previous results obtained by Quinn et al (38), where an increase in carbapenem resistance was observed when cells were exposed to 0.2 % HSA. As expected, since the AB5075 strain is already a highly carbapenem-resistant strain, additional increases in carbapenem MICs were not were observed (Table S2).

To expand the studies on the effect of HSA in the expression of carbapenem-resistance genes, three additional carbapenem-resistant strains, AMA16, ABUH 702, and AB0057, isolated from blood cultures were tested. These strains belong to three different clonal lineages and carry different carbapenem resistance mechanisms (Table S3) (42-45). The results showed that addition of 3.5% HSA to the culture medium of *A. baumannii* AMA16, ABUH702 and AB0057 was associated with an increase (4.1-, 1.6- and 14.8-fold, respectively) in the levels of *bla*_OXA-51-like_ mRNA (Fig S2). In the same conditions, transcription levels of *carO* decreased by 0.4-, 0.6- and 0.6-fold, respectively (Fig S2). Other genes that changed expression levels in the presence of 3.5% HSA were the *A. baumannii bla*_OXA-23_ (increased 1.7-fold), the *A. baumannii* ABUH702 IS*Aba1*transposase (increased 2.5-fold) (Fig. S2), and the *A. baumannii* AMA16 *bla*_NDM-1_ (decreased 0.4-fold) (Fig S2). These results are in keeping with those obtained with *A. baumannii* A118 and AB5075, suggesting that HSA could be a factor in the carbapenem failure to treat infections caused by numerous *A. baumannii* strains. Unfortunately, carbapenem resistance is widespsread in *A. baumannii*, for example a review of data analysis of the 2010-2014 and 2015-2019 cohorts of *A. baumannii* blood stream infections at Louis Stokes Cleveland Department of Veterans Affairs Medical Center, showed carbapenem resistance in 35% and 17% (Personal Communication).

When drugs are administered, their distribution is strongly controlled by HSA as most drugs circulate in plasma and reach the target tissues bound to this protein. Binding of a drug to albumin results in increased solubility in plasma, decreased toxicity, and protection against oxidation (46). Studies on the interaction between meropenem and HSA showed changes in the secondary structure and cleavage of the protein, which help the transportation and distribution of drug in blood plasma (46). Other studies also suggested that neither the full-length nor the tertiary structure of HSA are required for a given functionality, e.g., its effect on natural transformation in *A. baumannii* (13). We hypothesize that after binding HSA, carbapenems induce fragmentation and the peptides produced have the property to trigger expression of genes involved in natural competence or carbapenem resistance.

### 2.3. Pooled normal human serum affects gene expression of natural transformation associated genes as well as carbapenem-resistance genes

Building on our previous observations that human serum (HS), whose main component is HSA, enhances the competence of *A. baumannii* (13, 18), we tested the HS effect on the expression of competence and carbapenem-resistance genes. The expression of competence and type IV pilus associated genes were up-regulated (Fig. 3A). For *pilA* gene, a 20.5- and 9.1-fold increase was observed in the presence of HS for A118 and AB5075, respectively. The *pilQ* expression level increased by 9- and 17-fold under HS exposure for A118 and AB5075, respectively. Also, the expression level of *pilT* was 39- and 28-fold higher in the HS condition for A118 and AB5075, respectively. For *comEA* gene, we observed an increase of 2.4- and 10.1-fold for A118 and AB5075, respectively (Fig. 3A). Consistently, the transcripts levels of *comEC* were 9.5-fold (A118) and 2.5-fold (AB5075), while *comF* transcripts level values were increased by 28- and 5.4-fold, respectively for A118 and AB5075. Finally, *drpA* expression level was increased by 7- and 9-fold under HS conditions for A118 and AB5075, respectively (Fig. 3A). The results for the expression of competence-associated genes upon exposure to HS are in accordance with the induction of the expression of these genes in the presence of 3.5% HSA.

**Figure 3.**
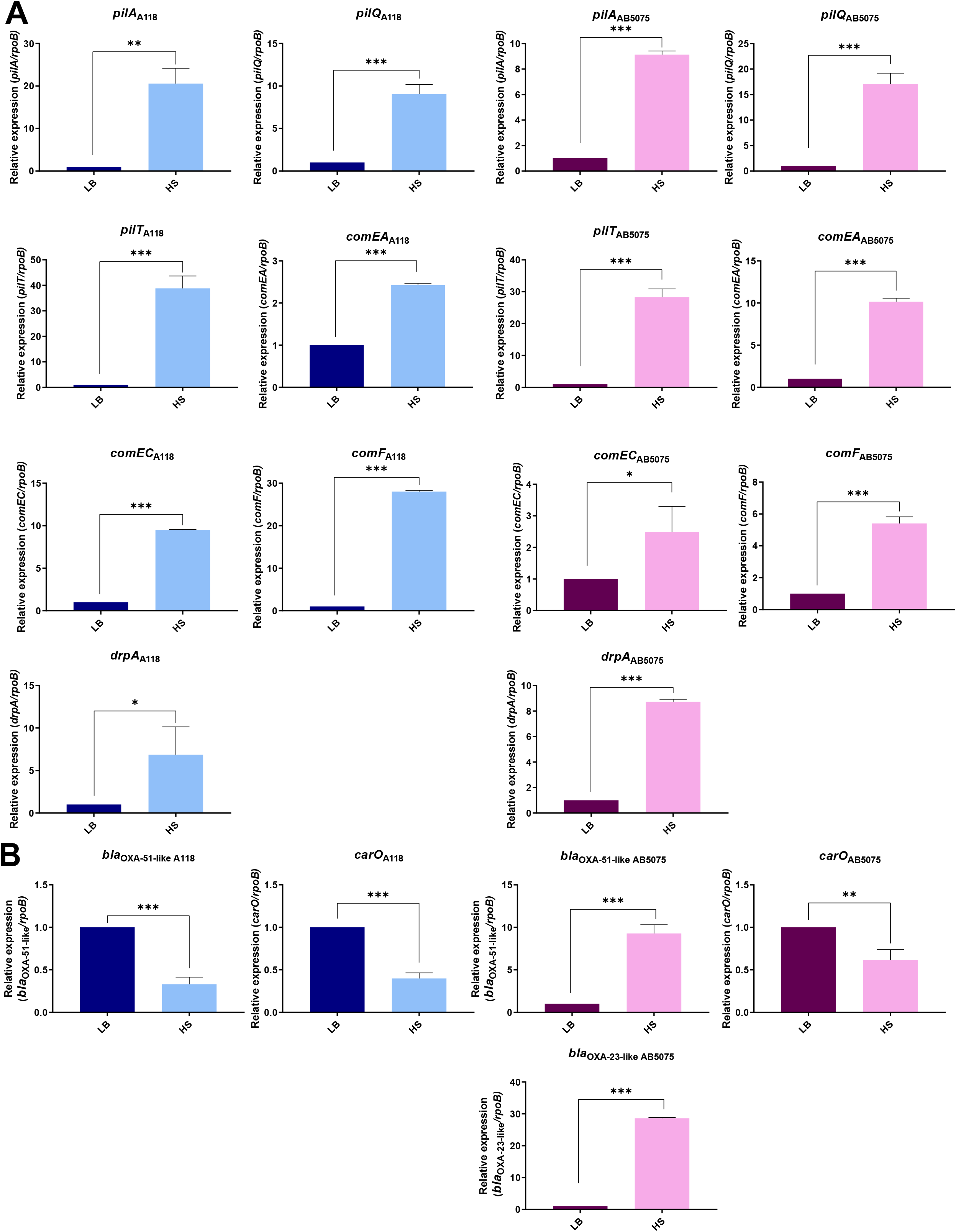
A) qRT-PCR of A118 and AB5075 strains genes associated with competence and type IV pilus, *pilA, pilQ, pilT, comEA, comEC, comF* and *dprA* in the presence of HS. B) qRT-PCR of carbapenem resistance related genes (*bla*_OXA-51-like_, *bla*_OXA-23-like_ and *carO*) for the *A. baumannii* A118 and AB5075 strains in the presence of HS. Bacterial cells grown in LB or HS were used. Fold changes were calculated using double ΔCt analysis. At least three independent samples were used, and three technical replicates were performed from each sample. Statistical significance (*P* < 0.05) was determined by *t* test, one asterisks: *P* < 0.05; two asterisks: *P* < 0.01 and three asterisks: *P* < 0.001.

To further explore the role of HSA as the main protein component of HS, we evaluated the impact of HS on the expression of carbapenem-resistance gene. qRT-PCR of select genes from A118 and AB5075 grown in HS was performed. Figure 3B reveals the levels of expression of carbapenem-resistance genes under HS treatment. We observed down-regulation of the transcript levels of *bla*_OXA-51-like_ for A118 strain in the presence of HS (0.33-fold decreased) (Fig. 3B), while an increase in the expression of *bla*_OXA-51-like_ and *bla*_OXA-23-like_ for AB5075 strains. The *carO* expression levels for both strains were down-regulated in both treatments with respect to the control (Fig. 3B), as previously observed for the 3.5 % HSA treatment. Carbapenem MIC results for both strains showed an increase in resistance for doripenem and ertapenem for A118, and imipenem and meropenem for AB5075 in the presence of HS (Table S4, Fig S3). The increase in imipenem and meropenem MICs was more than >3 doubling dilutions for AB5075. This effect was not previously observed with 3.5 % HSA; however, it may be the result of the increased expression of *bla*_OXA-51-like_ in this condition compared to the exposure to HSA (2,97-fold-change increase). Overall, these results suggest that HSA and HS effect the expression of genes involved in carbapenem-resistance, which is manifested in the observed phenotypes, contributing to the extraordinary capacity of *A. baumannii* to respond to human products by changing the expression of genes.

### 2.4. Natural transformation and the expression of competence and carbapenem-resistance genes are boosted in presence the of HSA and meropenem

Since HSA and carbapenems regulate a common group of genes, we tested the possibility that they can exert synergistic effects. *A. baumannii* A118 and AB5075 cells were cultured in medium supplemented with two-times sub-MICs (0.25X MIC) of meropenem (0.063 μg/mL and 8 μg/mL, respectively) with or without the addition of 3.5% HSA. While addition of meropenem increased transformation frequencies by 2-fold (Fig. 4A), the presence of both meropenem and HSA resulted in an increase of 44- (A118) and 8.7-fold (AB5075) (Fig. 4A). A comparison of these increases in transformation efficiencies with those measured in the sole presence of HSA showed divergent results. The combined effect of meropenem and HSA on transformation efficiency in *A. baumannii* A118 was synergistic (Fig. 4A). Conversely, addition of meropenem to HSA did not enhance transformation efficiency in *A. baumannii* AB5075 (Fig. S4A). Since this latter strain is resistant to meropenem, the lower stress caused by the presence of the antibiotic may be the reason behind the lack of synergistic effect.

**Figure 4.**
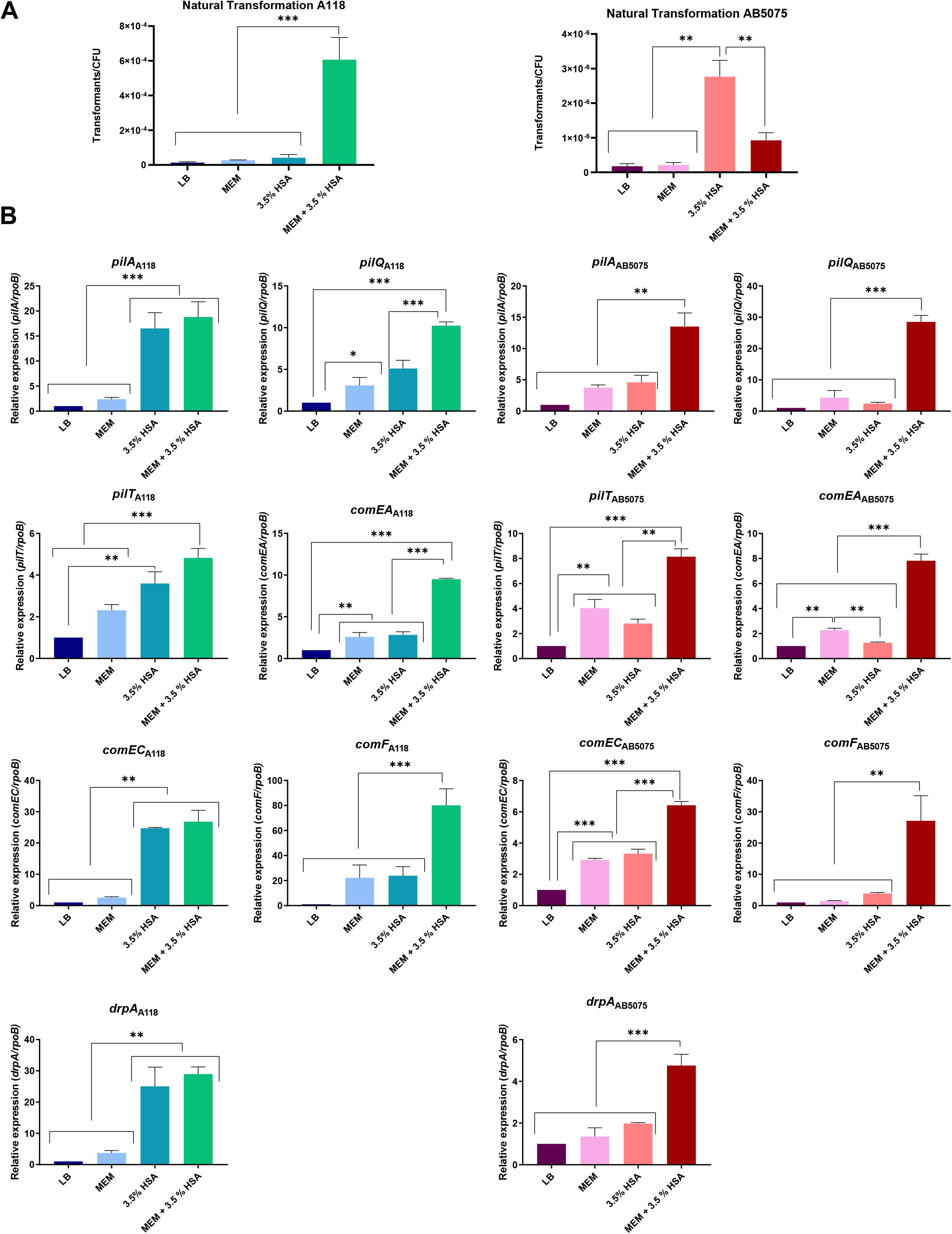
A) Natural transformation frequencies in LB broth, LB broth plus sub-MIC of meropenem (MEM), 3.5 % HSA or sub-MIC of MEM plus 3.5 % HSA for A118 and AB5075 *A. baumannii* strains. At least three independent replicates were performed and *P-value* < 0.05 was considered significant (ANOVA followed by Tukey’s multiple-comparison test). B) qRT-PCR of A118 and AB5075 strains genes associated with competence and type IV pilus, *pilA, pilQ, pilT, comEA, comEC, comF* and *dprA* in the presence of sub-MIC of MEM, 3.5 % HSA or sub-MIC of MEM plus 3.5% HSA. Fold changes were calculated using double ΔCt analysis. At least three independent samples were used, and three technical replicates were performed from each sample. Statistical significance (*P* < 0.05) was determined by ANOVA followed by Tukey’s multiple-comparison test; one asterisks: *P* < 0.05; two asterisks: *P* < 0.01 and three asterisks: *P* < 0.001.

The expression levels of competence and type IV pilus associated genes in the presence of sub-MIC concentrations of meropenem alone and in combination with 3.5 % HSA are shown in Figure 3B. Expression of all seven genes tested was enhanced in the presence of meropenem or 3.5% HSA in *A. baumannii* strains A118 and AB5075 (Fig. 4B). The increase in expression levels were comparable in the case of all seven *A. baumannii* AB5075 genes but in the case of strain A118, expression of *pilA, pilT, comEC*, and *drpA* was enhanced more in the presence of HSA (Fig. 4B). With the exception of these four genes whose levels did not change significantly in the presence of HSA or the combination of meropenem and HSA, all other genes were expressed at much higher levels in the presence of the combination than when the cells grew in the presence of one of the components (Fig. 4B).

Addition of a sub-MIC MEM concentration or 3.5 % HSA induced a modest increase in levels of expression of *bla*_OXA-51-like_ in *A. baumannii* A118 and AB5075 (Fig. 5). However, the presence of both compounds in the culture medium produced a synergistic effect, the expression levels of *bla*_OXA-51-like_ were dramatically increased (Fig. 5). The expression levels of the *A. baumannii* AB5075 *bla*_OXA-23-like_ followed a similar pattern but the enhancement produced by addition of 3.5 % HSA was more substantial than that observed for *bla*_OXA-51-like_ (Fig. 5). In the case of the *carO* gene, as expected, the expression levels in both strains were reduced in the presence of sub-MIC meropenem concentration or 3.5 % HSA (Fig. 5). The combination of both compounds produced a slight reduction that was not enough to qualify as synergistic (Fig. 5). These results are in agreement with a previous study conducted by Qin et al. (2018) that analyzed the transcriptomic response of three *A. baumannii* strains with different resistance and growth profiles showing that addition of sub-MIC concentrations of meropenem to the culture medium were associated with increased expression of β-lactamase genes in all strains tested (47). Thus, our results agree with this previous observation and contribute with additional information demonstrating that HSA further alters the transcriptional levels of carbapenem-resistance genes.

**Figure 5.**
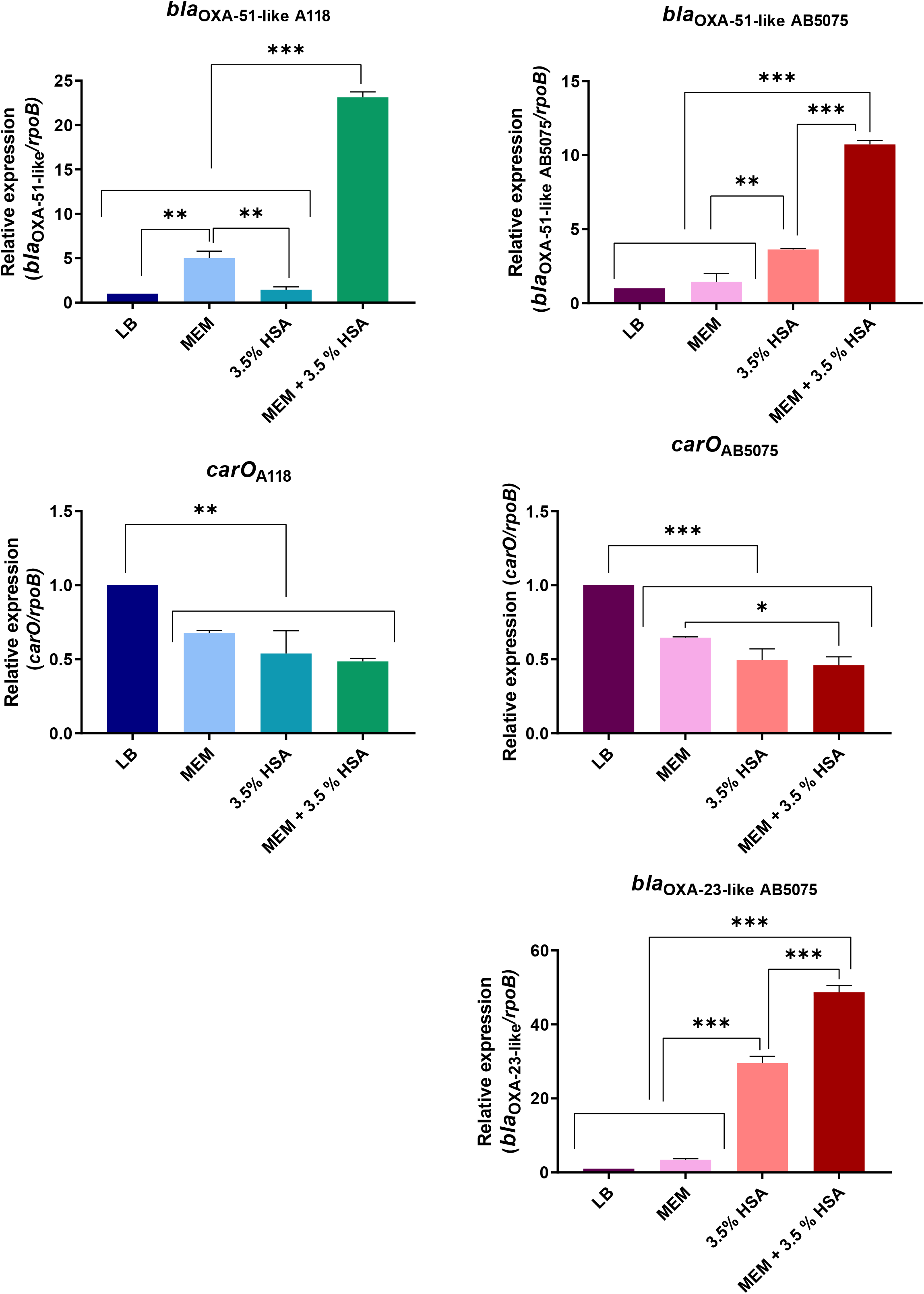
qRT-PCR of carbapenem-resistance genes (*bla*_OXA-51-like_, *bla*_OXA-23-like_ and *carO*) for the *A. baumannii* A118 and AB5075 strains in the presence of sub-MIC (0.25X MIC) of meropenem (MEM), 3.5% HSA and sub-MIC of MEM plus 3.5% HSA. Fold changes were calculated using double ΔCt analysis. At least three independent samples were used, and three technical replicates were performed from each sample. Statistical significance (*P* < 0.05) was determined by ANOVA followed by Tukey’s multiple-comparison test, one asterisks: *P* < 0.05; two asterisks: *P* < 0.01 and three asterisks: *P* < 0.001.

## 4. CONCLUSIONS

Bloodstream infections, associated with high mortality rates among hospitalized patients, are a significant threat to the public health system (4). Most importantly, high levels of multidrug resistance in *A. baumannii* significantly impact treatment options for those that develop bacteremia (48). Our study reveals an important interplay that is likely to occur during bloodstream *A. baumannii* infections that are treated with carbapenems. Specifically, we identified that HSA, the main protein component in HS, triggers natural transformation and also affects the expression level of antibiotic resistance genes. Furthermore, when HSA is combined with carbapenems, commonly used to treat bacterial infections such as bacteremia; the capacity of *A. baumannii* to acquire DNA and modified the expression of carbapenem-resistance genes.

In sum, the results described in this article show that *A. baumannii* responds to conditions that resemble those found in the human body in untreated or carbapenem-treated infection, augmenting its transformability and modifying the expression of resistance genes. The *A. baumannii’s* ability to sense and respond to human products or antibiotics modulating evolutive capacities and antimicrobial resistance levels contributes to the success of this bacterium as a threat to human health.

## 3. MATERIAL AND METHODS

### 3.1. Bacterial strains and plasmid

The model *A. baumannii* strain A118 and AB5075, which show a different degree of susceptibility and virulence, were used (8, 35, 36, 49, 50). AMA 16 strain (NDM-1 positive strains), ABUH702 (carbapenem resistance due to increase expression of *bla*_OXA-66_ by IS*Aba1*), and AB0057 (OXA-23 positive strain) were also used (42-45). The pMBLe-OA-ArK plasmid, including the apramycin (ArkR), was used for transformation assays (51). Plasmid DNA was obtained from *Escherichia coli* TOP10 cells harboring pMBLe-OA-ArK using QIAprep Spin Miniprep Kit following manufacturer instructions (Qiagen Germantown, MD, USA).

### 3.2. Natural transformation assays

Natural transformation assays were adapted from the previously described protocol used by Vesel et al. Briefly, *A. baumannii* cells were grown overnight in LB medium and diluted 1:100 into 2 mL of LB medium with or without 3.5% HSA, or 2 times sub-MIC of meropenem (0.0625 ug/ml and 8 ug/ml for A118 and AB5075, respectively), or 3.5% HSA plus 2 times sub-MIC of meropenem. The cultures were incubated aerobically at 37^○^C until they reached an OD600 of approximately 0.65. After, 20 μL of the bacterial culture was mixed with 1 μg of plasmid DNA (pMBLe-OA-ArK) and the mixture was spotted onto motility plates. After 2 h of incubation at 37 C, the cells were scraped off from the plate and resuspended in an eppendorf tube containing 200 μL of LB medium. Transformation events were scored by counting apramycin colonies, while total colony forming units (CFUs) was assessed by plating serial dilutions on LB agar plates. Negative controls with no DNA addition were included in every tested condition. All experiments were performed in triplicate and statistical analysis was performed. Transformation events were scored as mentioned above.

### 3.3. RNA extraction and quantitative reverse transcription polymerase chain reaction (qRT-PCR)

Overnight cultures of A118 and AB5075 were then diluted 1:10 in fresh LB medium or LB medium supplemented with 3.5% HSA, or pooled normal human serum, or 2 times sub-MIC of meropenem (0.0625 ug/ml and 8 ug/ml for A118 and AB5075, respectively), or 3.5 % HSA + 2 times sub-MIC of meropenem and incubated with agitation for 5 h at 37°C. In addition, AMA16, ABUH702 and AB0057 cultured in LB medium or LB medium supplemented with 3.5% HSA were also evaluated the expression of carbapenem-resistance genes. Pure HSA (Sigma-Aldrich, St. Louis, MO, USA) and pooled normal human serum from a certified vendor (Innovative Research Inc, Novi, MI, USA) were used in the cultures.

RNA was extracted from each strain using the Direct-zol RNA Kit (Zymo Research, Irvine, CA, USA) following manufacturer’s instructions. Total RNA extractions were performed in three biological replicates for each condition. The extracted and DNase-treated was used to synthesized cDNA using the manufacturer protocol provided within the iScript™ Reverse Transcription Supermix for qPCR (Bio-Rad, Hercules, CA, United States). The cDNA concentrations were adjusted to 50 ng/μl and qPCR was conducted using the qPCRBIO SyGreen Blue Mix Lo-ROX following manufacturer’s protocol (PCR Biosystems, Wayne, Pennsylvania, USA). At least three biological replicates of cDNA were used in triplets and were run using the CFX96 Touch™ Real-Time PCR Detection System (Bio-Rad, Hercules, CA, USA).

Transcriptional levels of each sample were normalized to the transcriptional level of *rpoB*. The relative quantification of gene expression was performed using the comparative threshold method 2^-ΔΔCt^. The ratios obtained after normalization were expressed as folds of change compared with cDNA samples isolated from bacteria cultures on LB. Asterisks indicate significant differences as determined by ANOVA followed by Tukey’s multiple comparison test (*P* < 0.05), using GraphPad Prism (GraphPad software, San Diego, CA, United States).

### 3.4. Antimicrobial susceptibility testing

As previously described (52), Antibiotic susceptibility assays were performed following with the Procedures recommended by the CLSI. After OD adjustment, 100 μl of cells grown in LB or LB plus 3.5 % or pooled normal human serum were inoculated on Mueller-Hinton agar plates. Antimicrobial commercial E-strips (Liofilchem S.r.l., Italy) for meropenem (MER), imipenem (IPM), doripenem (DOR), and ertapenem (ERT) were used. Mueller-Hinton agar plates were incubated at 37°C for 18 h. CLSI breakpoints were used for interpretation (53). (53). MICs for meropenem (MEM), imipenem (IMP), doripenem (DOR), and ertapenem (ERT) were carried out as previously described (38).

### 3.5. Statistical Analysis

All experiments were performed at least in three technical and biological replicates. Data were expressed as means ± standard deviation. Statistical analysis using *t* test or ANOVA followed by Tukey’s multiple comparison test were performed using GraphPad Prism (GraphPad software, San Diego, CA, USA), and a *P* value < 0.05 was considered statistically significant.

## Supplementary Materials

**Table S1**. Heatmap showing the differential expression of competence and type IV pilus associated genes pilus associated in *A. baumannii* A118 and AB5075 strains under 0.2% HSA, 4% HPF and 4% CSF induction (11, 35-37). (Do we add our references?).

**Table S2**. Minimal Inhibitory Concentrations of *A. baumannii* strains A118 and AB5075 grew in LB or LB plus 3.5 % HSA.

**Table S3**. Characteristics of three additional carbapenem-resistant strains isolated from blood samples included in the present study.

**Table S4**. Carbapenems MICs of A118 or AB5075 grew in LB or Human Serum.

**Figure S1**. Heatmap outlining the differential expression of genes associated with type IV pilus in the presence of 0.2% HSA, 4% HPF and 4% CSF in *A. baumannii* A118 and AB5075 strains. The asterisks represent significant DEGs (adjusted *P*-value < 0.05 with log2fold change > 1).

**Figure S2**. Determination of relative expression levels of carbapenem-associated genes in carbapenem-resistant strains isolated from blood samples upon LB and LB plus 3.5 % exposure. qPCR was used to determine the expression levels and the fold changes were using double ΔCt analysis. At least two independent samples were used, and three technical replicates were performed from each sample. Statistical significance (*P* < 0.05) was determined by ANOVA followed by *t* test; one asterisks: *P* < 0.05; two asterisks: *P* < 0.01 and three asterisks: *P* < 0.001.

**Figure S3**. A) Strain A118 grew in LB broth or HS were used to performed doripenem (DOR) and ertapenem (ERT) susceptibility assays. B) Strain AB5075 grew in LB broth or HS were used to performed meropenem (MEM) and imipenem (IMP) susceptibility. Minimum inhibitory concentration (MIC) was performed by E-test (Liofilchem, Italy) following CLSI recommendations.

## Author Contributions

C.P., C.L., M.R.T., T.S., R.A.B, and M.S.R. conceived the study and designed the experiments. C.P., C.L., M.R.T., T.S., B.N., and M.S.R. performed the experiments and genomics and bioinformatics analyses. C.P., C.L., M.R.T., T.S., G.M.T., F.P., K.M.P., R.A.B., M.E.T. and M.S.R. analyzed the data and interpreted the results. K.P.W., R.A.B., M.E.T. and M.S.R. contributed reagents/materials/analysis tools. M.R.T., T.S., F.P., K.M.P., R.A.B.,M.E.T. and M.S.R. wrote and revised the manuscript. All authors read and approved the final manuscript.

## Funding

The authors’ work was supported by NIH SC3GM125556 to MSR, R01AI100560 to RAB, R01AI063517, R01AI072219 to RAB, and 2R15 AI047115 to MET. This study was supported in part by funds and/or facilities provided by the Cleveland Department of Veterans Affairs, Award Number 1I01BX001974 to RAB from the Biomedical Laboratory Research & Development Service of the VA Office of Research and Development and the Geriatric Research Education and Clinical Center VISN 10 to RAB. The content is solely the responsibility of the authors and does not necessarily represent the official views of the National Institutes of Health or the Department of Veterans Affairs. MRT and TS are recipient of a postdoctoral fellowship from CONICET.

## Conflicts of Interest

The authors declare no conflict of interest.

## REFERENCES

1. CDC. 2019. Antibiotic resistance threats in the United States. Centers for Disease Control, Atlanta, GA.

2. Isler B, Doi Y, Bonomo RA, Paterson DL. 2019. New treatment options against carbapenem-resistant Acinetobacter baumannii infections. Antimicrob Agents Chemother 63.

3. Eichenberger EM, Thaden JT. 2019. Epidemiology and Mechanisms of Resistance of Extensively Drug Resistant Gram-Negative Bacteria. Antibiotics (Basel) 8.

4. Holmes CL, Anderson MT, Mobley HLT, Bachman MA. 2021. Pathogenesis of Gram-Negative Bacteremia. Clin Microbiol Rev 34.

5. Garcia-Ortega L, Arch O, Perez-Canosa C, Lupion C, Gonzalez C, Rodriguez-Bano J, Spanish Study Group of Nosocomial I. 2011. Control measures for Acinetobacter baumannii: a survey of Spanish hospitals. Enferm Infecc Microbiol Clin 29:36–8.

6. Harding CM, Tracy EN, Carruthers MD, Rather PN, Actis LA, Munson RS, Jr. 2013. Acinetobacter baumannii strain M2 produces type IV pili which play a role in natural transformation and twitching motility but not surface-associated motility. MBio 4.

7. Pour NK, Dusane DH, Dhakephalkar PK, Zamin FR, Zinjarde SS, Chopade BA. 2011. Biofilm formation by Acinetobacter baumannii strains isolated from urinary tract infection and urinary catheters. FEMS Immunol Med Microbiol 62:328–38.

8. Ramirez MS, Don M, Merkier AK, Bistue AJ, Zorreguieta A, Centron D, Tolmasky ME. 2010. Naturally competent Acinetobacter baumannii clinical isolate as a convenient model for genetic studies. J Clin Microbiol 48:1488–90.

9. Ramirez MS, Merkier AK, Quiroga MP, Centron D. 2012. Acinetobacter baumannii is able to gain and maintain a plasmid harbouring In35 found in Enterobacteriaceae isolates from Argentina. Curr Microbiol 64:211–3.

10. Wilharm G, Piesker J, Laue M, Skiebe E. 2013. DNA uptake by the nosocomial pathogen Acinetobacter baumannii occurs during movement along wet surfaces. J Bacteriol 195:4146–53.

11. Quinn B, Traglia GM, Nguyen M, Martinez J, Liu C, Fernandez JS, Ramirez MS. 2018. Effect of Host Human Products on Natural Transformation in Acinetobacter baumannii. Curr Microbiol.

12. Quinn B, Martinez J, Liu C, Nguyen M, Ramirez MS. 2018. The effect of sub-inhibitory concentrations of antibiotics on natural transformation in Acinetobacter baumannii. Int J Antimicrob Agents 51:809–810.

13. Traglia GM, Quinn B, Schramm ST, Soler-Bistue A, Ramirez MS. 2016. Serum Albumin and Ca2+ Are Natural Competence Inducers in the Human Pathogen Acinetobacter baumannii. Antimicrob Agents Chemother 60:4920–9.

14. Godeux AS, Svedholm E, Lupo A, Haenni M, Venner S, Laaberki MH, Charpentier X. 2020. Scarless Removal of Large Resistance Island AbaR Results in Antibiotic Susceptibility and Increased Natural Transformability in Acinetobacter baumannii. Antimicrob Agents Chemother 64.

15. Domingues S, Rosario N, Candido A, Neto D, Nielsen KM, Da Silva GJ. 2019. Competence for Natural Transformation Is Common among Clinical Strains of Resistant Acinetobacter spp. Microorganisms 7.

16. Domingues S, Rosario N, Ben Cheikh H, Da Silva GJ. 2018. ISAba1 and Tn6168 acquisition by natural transformation leads to third-generation cephalosporins resistance in Acinetobacter baumannii. Infect Genet Evol 63:13–16.

17. Hu Y, He L, Tao X, Meng F, Zhang J. 2019. High DNA Uptake Capacity of International Clone II Acinetobacter baumannii Detected by a Novel Planktonic Natural Transformation Assay. Front Microbiol 10:2165.

18. Martinez J, Liu C, Rodman N, Fernandez JS, Barberis C, Sieira R, Perez F, Bonomo RA, Ramirez MS. 2018. Human fluids alter DNA-acquisition in Acinetobacter baumannii. Diagn Microbiol Infect Dis.

19. Gerischer U, Ornston LN. 2001. Dependence of linkage of alleles on their physical distance in natural transformation of Acinetobacter sp. strain ADP1. Arch Microbiol 176:465–9.

20. Griffith F. 1928. The Significance of Pneumococcal Types. J Hyg (Lond) 27:113–59.

21. Hahn J, Maier B, Haijema BJ, Sheetz M, Dubnau D. 2005. Transformation proteins and DNA uptake localize to the cell poles in Bacillus subtilis. Cell 122:59–71.

22. Johnston C, Martin B, Fichant G, Polard P, Claverys JP. 2014. Bacterial transformation: distribution, shared mechanisms and divergent control. Nat Rev Microbiol 12:181–96.

23. Lorenz MG, Wackernagel W. 1994. Bacterial gene transfer by natural genetic transformation in the environment. Microbiol Rev 58:563–602.

24. Dubnau D, Blokesch M. 2019. Mechanisms of DNA Uptake by Naturally Competent Bacteria. Annu Rev Genet 53:217–237.

25. Meibom KL, Blokesch M, Dolganov NA, Wu CY, Schoolnik GK. 2005. Chitin induces natural competence in Vibrio cholerae. Science 310:1824–7.

26. Ellison CK, Dalia TN, Vidal Ceballos A, Wang JC, Biais N, Brun YV, Dalia AB. 2018. Retraction of DNA-bound type IV competence pili initiates DNA uptake during natural transformation in Vibrio cholerae. Nat Microbiol 3:773–780.

27. Charpentier X, Polard P, Claverys JP. 2012. Induction of competence for genetic transformation by antibiotics: convergent evolution of stress responses in distant bacterial species lacking SOS? Curr Opin Microbiol 15:570–6.

28. Prudhomme M, Attaiech L, Sanchez G, Martin B, Claverys JP. 2006. Antibiotic stress induces genetic transformability in the human pathogen Streptococcus pneumoniae. Science 313:89–92.

29. Ramirez MS, Bonomo RA, Tolmasky ME. 2020. Carbapenemases: Transforming Acinetobacter baumannii into a Yet More Dangerous Menace. Biomolecules 10.

30. Vesel N, Blokesch M. 2021. Pilus production in Acinetobacter baumannii is growth phase dependent and essential for natural transformation. J Bacteriol.

31. Traglia GM, Chua K, Centron D, Tolmasky ME, Ramirez MS. 2014. Whole-genome sequence analysis of the naturally competent Acinetobacter baumannii clinical isolate A118. Genome Biol Evol 6:2235–9.

32. Boyd JM, Koga T, Lory S. 1994. Identification and characterization of PilS, an essential regulator of pilin expression in Pseudomonas aeruginosa. Mol Gen Genet 243:565–74.

33. Ishimoto KS, Lory S. 1992. Identification of pilR, which encodes a transcriptional activator of the Pseudomonas aeruginosa pilin gene. J Bacteriol 174:3514–21.

34. Hobbs M, Collie ES, Free PD, Livingston SP, Mattick JS. 1993. PilS and PilR, a two-component transcriptional regulatory system controlling expression of type 4 fimbriae in Pseudomonas aeruginosa. Mol Microbiol 7:669–82.

35. Martinez J, Razo-Gutierrez C, Le C, Courville R, Pimentel C, Liu C, Fung SE, Tuttobene MR, Phan K, Vila AJ, Shahrestani P, Jimenez V, Tolmasky ME, Becka SA, Papp-Wallace KM, Bonomo RA, Soler-Bistue A, Sieira R, Ramirez MS. 2021. Cerebrospinal fluid (CSF) augments metabolism and virulence expression factors in Acinetobacter baumannii. Sci Rep 11:4737.

36. Rodman Nyah MJ, Fung Sammie, Nakanouchi Jun, Myers Amber L., Harris Caitlin M., Dang Emily, Fernandez Jennifer S., Liu Christine, Mendoza Anthony M., Jimenez Veronica, Nikolaidis Nikolas, Brennan Catherine A., Bonomo Robert A., Sieira Rodrigo, Ramirez Maria Soledad. 2019. Human Pleural Fluid Elicits Pyruvate and Phenylalanine Metabolism in Acinetobacter baumannii to Enhance Cytotoxicity and Immune Evasion. Frontiers in Microbiology 10:1581.

37. Martinez J, Fernandez JS, Liu C, Hoard A, Mendoza A, Nakanouchi J, Rodman N, Courville R, Tuttobene MR, Lopez C, Gonzalez LJ, Shahrestani P, Papp-Wallace KM, Vila AJ, Tolmasky ME, Bonomo RA, Sieira R, Ramirez MS. 2019. Human pleural fluid triggers global changes in the transcriptional landscape of Acinetobacter baumannii as an adaptive response to stress. Sci Rep 9:17251.

38. Quinn B, Rodman N, Jara E, Fernandez JS, Martinez J, Traglia GM, Montana S, Cantera V, Place K, Bonomo RA, Iriarte A, Ramirez MS. 2018. Human serum albumin alters specific genes that can play a role in survival and persistence in Acinetobacter baumannii. Sci Rep 8:14741.

39. Fanali G, di Masi A, Trezza V, Marino M, Fasano M, Ascenzi P. 2012. Human serum albumin: from bench to bedside. Mol Aspects Med 33:209–90.

40. Wong D, Nielsen TB, Bonomo RA, Pantapalangkoor P, Luna B, Spellberg B. 2017. Clinical and pathophysiological overview of Acinetobacter infections: a century of challenges. Clin Microbiol Rev 30:409–447.

41. Lomaestro BM, Drusano GL. 2005. Pharmacodynamic evaluation of extending the administration time of meropenem using a Monte Carlo simulation. Antimicrob Agents Chemother 49:461–3.

42. Adams MD, Pasteran F, Traglia GM, Martinez J, Huang F, Liu C, Fernandez JS, Lopez C, Gonzalez LJ, Albornoz E, Corso A, Vila AJ, Bonomo RA, Ramirez MS. 2020. Distinct mechanisms of dissemination of NDM-1 metallo-beta-lactamase in Acinetobacter spp. in Argentina. Antimicrob Agents Chemother.

43. Rodgers D, Pasteran F, Calderon M, Jaber S, Traglia GM, Albornoz E, Corso A, Vila AJ, Bonomo RA, Adams MD, Ramirez MS. 2020. Characterisation of ST25 NDM-1-producing Acinetobacter spp. strains leading the increase in NDM-1 emergence in Argentina. J Glob Antimicrob Resist 23:108–110.

44. Adams MD, Wright MS, Karichu JK, Venepally P, Fouts DE, Chan AP, Richter SS, Jacobs MR, Bonomo RA. 2019. Rapid Replacement of Acinetobacter baumannii Strains Accompanied by Changes in Lipooligosaccharide Loci and Resistance Gene Repertoire. mBio 10.

45. Adams MD, Goglin K, Molyneaux N, Hujer KM, Lavender H, Jamison JJ, MacDonald IJ, Martin KM, Russo T, Campagnari AA, Hujer AM, Bonomo RA, Gill SR. 2008. Comparative genome sequence analysis of multidrug-resistant Acinetobacter baumannii. J Bacteriol 190:8053–8064.

46. Abdo Esmail SA, Shamsi M, Al-Asbahy WM. 2019. Interaction and photo-induced cleavage studies of meropenem drug with human serum albumin using spectroscopic and molecular docking investigations. J Biomol Struct Dyn 37:3282–3289.

47. Qin H, Lo NW, Loo JF, Lin X, Yim AK, Tsui SK, Lau TC, Ip M, Chan TF. 2018. Comparative transcriptomics of multidrug-resistant Acinetobacter baumannii in response to antibiotic treatments. Sci Rep 8:3515.

48. Diekema DJ, Hsueh PR, Mendes RE, Pfaller MA, Rolston KV, Sader HS, Jones RN. 2019. The Microbiology of Bloodstream Infection: 20-Year Trends from the SENTRY Antimicrobial Surveillance Program. Antimicrob Agents Chemother 63.

49. Jacobs AC, Thompson MG, Black CC, Kessler JL, Clark LP, McQueary CN, Gancz HY, Corey BW, Moon JK, Si Y, Owen MT, Hallock JD, Kwak YI, Summers A, Li CZ, Rasko DA, Penwell WF, Honnold CL, Wise MC, Waterman PE, Lesho EP, Stewart RL, Actis LA, Palys TJ, Craft DW, Zurawski DV. 2014. AB5075, a Highly Virulent Isolate of Acinetobacter baumannii, as a Model Strain for the Evaluation of Pathogenesis and Antimicrobial Treatments. MBio 5:e01076–14.

50. Ramirez MS, Penwell WF, Traglia GM, Zimbler DL, Gaddy JA, Nikolaidis N, Arivett BA, Adams MD, Bonomo RA, Actis LA, Tolmasky ME. 2019. Identification of Potential Virulence Factors in the Model Strain Acinetobacter baumannii A118. Front Microbiol 10:1599.

51. Huang F, Fitchett N, Razo-Gutierrez C, Le C, Martinez J, Ra G, Lopez C, Gonzalez LJ, Sieira R, Vila AJ, Bonomo RA, Ramirez MS. 2020. The H-NS Regulator Plays a Role in the Stress Induced by Carbapenemase Expression in Acinetobacter baumannii. mSphere 5.

52. Ramirez MS, Traglia GM, Perez JF, Muller GL, Martinez F, Golic AE, Mussi MA. 2015. White and blue light induce reduction in susceptibility to minocycline and tigecycline in Acinetobacter sp. and other bacteria of clinical importance. J Med Microbiol.

53. (CLSI) CLSI. 2018. Methods for dilution antimicrobial susceptibility tests for bacteria that growth aerobically M7-A10, Ed. M7-A10. Clinical Lab Standards Institute.

